# RNA-seq of the medusavirus suggests remodeling of the host nuclear environment at an early infection stage

**DOI:** 10.1101/2021.04.10.439121

**Authors:** Ruixuan Zhang, Hisashi Endo, Masaharu Takemura, Hiroyuki Ogata

## Abstract

Nucleo–cytoplasmic large DNA viruses (NCLDVs) undergo a cytoplasmic or nucleo–cytoplasmic cycle, and the latter involves both nuclear and cytoplasmic compartments to proceed viral replication. Medusavirus, a recently isolated NCLDV, has a nucleo–cytoplasmic replication cycle in amoebas during which the host nuclear membrane apparently remains intact, a unique feature among amoeba–infecting giant viruses. The medusavirus genome lacks most transcription genes but encodes a full set of histone genes. To investigate the infection strategy, we performed a time–course RNA–seq experiment. All the viral genes were transcribed and classified into five temporal expression clusters. The immediate early genes (cluster 1, 42 genes) were mostly (83%) of unknown functions, frequently (95%) associated with a palindromic promoter–like motif, and enriched (45%) in putative nuclear–targeting genes. The later genes (clusters 2–5) were assigned to various functional categories. The viral linker histone H1 gene was in cluster 1, whereas the four core histone genes were in cluster 3, suggesting they had distinct roles during the course of the virus infection. The transcriptional profile of the host amoeba, *Acanthamoeba castellanii*, genes was greatly altered post–infection. Several encystment–related host genes showed increased representation levels at 48 hours post–infection, which is consistent with the previously reported amoeba encystment upon medusavirus infection. Overall, the transcriptional landscape during the course of medusavirus infection suggests that the virus modifies the host nuclear environment immediately after the initiation of infection. –

**Importance:** Medusavirus is an amoeba-infecting giant virus that was isolated from a hot spring in Japan. It belongs to the proposed family “Medusaviridae” in the phylum *Nucleocytoviricota*. Unlike other amoeba-infecting giant viruses, medusavirus initiates its DNA replication in the host nucleus without disrupting the nuclear membrane. Our RNA-seq analysis of its infection course uncovered ordered viral gene expression profiles. We identified temporal expression clusters of viral genes and associated putative promoter motifs. The subcellular localization prediction showed a clear spatiotemporal correlation between gene expression timing and localization of the encoded proteins. Notably, the immediate early expression cluster was enriched in genes targeting the nucleus, suggesting the priority of remodeling the host intra-nuclear environment during infection. The transcriptional profile of the amoeba genes was greatly altered post-infection. Notably, the expression of encystment-related genes increased 48 hours post-infection, suggesting that encystment may be an antiviral strategy of amoeba.

## Introduction

Giant viruses are characterized by their large viral particles and complex genomes, and are found worldwide (1–6). They are commonly referred to as nucleo-cytoplasmic large DNA viruses (NCLDVs) (7) and have been classified into the phylum *Nucleocytoviricota* (8). Phylogenetic analyses suggested that the diversification of giant viruses predated the emergence of modern eukaryotic lineages (9, 10), which revived the debate on their evolutionary origin (11, 12) and their relationship to the genesis of the eukaryotic nucleus (13). Genomic analysis revealed a large number of genes (referred to as orphan genes) that had no detectable homology to any known genes. The abundance of orphan genes or lineage-specific genes has been considered as evidence that supports the ongoing *de novo* creation of genes in these viruses (14, 15). In addition to the efforts to isolate new giant viruses, environmental genomics has revealed their cosmopolitan nature, extensive gene transfers with eukaryotes, and complex metabolic capabilities (16–18).

Medusavirus, a giant virus that infects the amoeba *Acanthamoeba castellanii*, was isolated from a hot spring in Japan (2) and, recently, a related virus medusavirus stheno was isolated from fresh water in Japan (19). These two viruses represent the proposed family “Medusaviridae” (2), which is distantly related to other giant virus families and forms an independent branch in the tree of the phylum *Nucleocytoviricota*. Medusavirus is unique among currently characterized amoeba-infecting giant viruses. Its DNA enters the host nucleus to initiate genome replication but the nuclear membrane remains intact during its replication cycle, and particle assembly and DNA packaging are carried out in the cytoplasm. All other amoeba-infecting giant viruses characterized to date carry out either a cytoplasmic replication by establishing cytoplasmic viral factories (e.g., mimiviruses (20), marseilleviruses (21), pithoviruses (5), cedratvirus (22), orpheovirus (23)) or a nucleo-cytoplasmic replication, like medusavirus, but with a degradation of the host nucleus (e.g., pandoraviruses (15), molliviruses (1)). For medusavirus, no visible cytoplasmic virus factory has been observed by transmission electron microscopy (2). Thus, it appears that the host nucleus is transformed into a virus factory, from which mature and immature medusavirus virions emerge. It has also been reported that some of the host amoeba cells display encystment upon medusavirus infection as early as 48 hours post-infection (hpi) (2). Medusavirus has a 381-kb genome that encodes 461 putative proteins; 86 (19%) have their closest homologs in *A. castellanii*, whereas 279 (61%) are orphan genes. Compared with other amoeba-infecting giant viruses, medusaviruses have fewer transcriptional and translational genes and have no genes that encode RNA polymerases and aminoacyl-tRNA synthetases, suggesting that medusaviruses are heavily reliant on the host machinery for transcription and translation. In contrast to their paucity in expression-related genes, medusavirus is unique among known viruses in encoding a complete set of histone domains, namely the core histones (H2A, H2B, H3, H4) and the linker histone H1. A virion proteomic study detected proteins encoded by the four core histone genes in medusavirus particles (2). Given these unique features, medusavirus is expected to have a characteristic infection strategy among known amoeba-infecting giant viruses. However, the dynamics of gene expression during the medusavirus infection cycle has not been reported so far.

Previous RNA-seq studies of giant viruses detected viral genes that were expressed in a coordinated manner during the viral infection. Viral genes that belong to different functional categories tend to show different expression patterns, and can be grouped as, for instance, “early”, “intermediate”, or “late”. Different viruses also have different gene expression programs; for example, the transcription order of informational genes (those involved in transcription, translation, and related processes) can differ among viruses. The expression of DNA replication genes (starting from 3 hpi) precedes the expression of transcription-related genes (6 hpi) in mimiviruses, whereas this order is reversed in a marseilleviruses (i.e., transcription-related genes from <1 hpi, DNA replication genes at 1–2 hpi) (24, 25). Putative promoter motifs that are associated with temporal expression groups have been identified in mimiviruses and marseilleviruses (24–27). The expression patterns of host genes during infection of giant viruses have been investigated by RNA-seq or proteomics (1, 25, 27).

We performed a time-series RNA-seq analysis of infected amoeba cells to investigate the transcriptional program and infection strategy of medusavirus. We report expression clusters of medusavirus genes, putative viral promoter motifs, as well as changes in host gene expression.

## Materials and Methods

### Amoeba culture, virus infection, and sequencing

*Acanthamoeba castellanii* strain Neff (ATCC 30010) cells were purchased from the American Type Culture Collection (ATCC, Manassas, VA, USA). The *A. castellanii* cells were cultured in eight 75-cm^2^ flasks with 25 mL of peptone-yeast-glucose (PYG) medium at 26°C for 1 hour, then infected with medusavirus at a multiplicity of infection of 2.88. After addition of medusavirus to 7 of the 8 flasks (one was the negative control), cells were harvested from each flask at 1, 2, 4, 8, 16, 24, and 48 hpi. Each cell pellet was washed with 1 mL of PBS by centrifugation (500 × g, 5 min at room temperature). Total RNA extraction was performed with an RNeasy Mini Kit (QIAGEN Inc., Japan) and quality checked by agarose gel electrophoresis. The extracted RNA was sent to Macrogen Corp. Japan for cDNA synthesis and library construction.

The cDNA synthesis and library construction were done using a TruSeq Stranded mRNA LT Sample Prep Kit (Illumina Inc., San Diego, CA, USA) following the manufacturer’s protocol. Briefly, the poly-A-containing mRNAs were purified using poly-T oligo attached magnetic beads. Then, the mRNA was fragmented using divalent cations under elevated temperature. First-strand cDNA was obtained using reverse transcriptase and random primers. After second-strand synthesis, the cDNAs were adenylated at their 3′ ends, and the adaptors were added. The DNA fragments were amplified by PCR and purified to create the final cDNA library. The RNA-seq was performed on a NovaSeq 6000 platform (Illumina Inc.).

### Read mapping and count normalization

The quality of the obtained reads was checked using the FastQC tool (http://www.bioinformatics.babraham.ac.uk/projects/fastqc/), which showed that the overall quality was above the threshold (quality threshold ≥20, no known adapters). Thus, we did no further trimming of the reads. The mRNA reads were mapped to a merged dataset composed of the nuclear genome of *A. castellanii* (GCF_000313135.1_Acastellanii.strNEFF_v1), the medusavirus genome (AP018495.1), and the mitochondria genome of *A. castellanii* (NC_001637.1) using HISAT2 (28) with maximum intron size of 1000. The number of reads mapped on each gene was calculated using HTSeq in union mode (29). The transcriptional activity of genes was estimated by Reads per Kilobases of transcript per Million mapped reads (RPKM) (Supplementary Material 1).

### Clustering

To discover the transcriptional patterns during medusavirus infection, we clustered the gene transcription profiles using the k-means method. We chose the library from 0–16 hpi to cluster the genes because viral DNA was observed in the cytoplasm at approximately 14 hpi and new virions were observed to be released, which indicated that a cycle of infection had ended (2). Genes with at least one mapped read across the 0–16 hpi libraries were included in the downstream analysis. To define the optimal number of clusters without the need for prior biological information, we used the R packages NbClust and clusterCrit, which use different clustering indices to estimate the quality of clusters (30, 31). For virus genes, most indices gave 5 as the optimal number of clusters, and for amoeba nuclear genes, most indices gave 2 as the optimal number of clusters. Therefore, we performed the k-means clustering with k = 5 and 2 for viral and amoeba nuclear genes, respectively (Supplementary Material 2).

### Subcellular localization prediction of viral genes

Subcellular localization prediction of medusavirus genes was done using the DeepLoc-1.0 server (32). The “plastid” category was removed, and the second-best prediction was chosen. We also predicted the subcellular localization tendency of other amoeba-infecting NCLDVs using the same method (Supplementary Material 3&4).

### Sequence motif analysis

MEME v5.1.1 was used for de novo motif prediction in the 5’ upstream sequence of medusavirus (33). We extracted 150-bp sequences upstream of the open reading frames. MEME was used in classic mode with motif width ranges of 8–10 bp and 6–15 bp to 8–25 bp, and “zero or more motifs in each intergenic region”. We selected the results in the 8–25-bp range because this was the only range in which the palindromic motif was found (Supplemental Material 4). We used the FIMO software tool (34) to scan the medusavirus genome for motifs that were predicted by MEME. The RNAMotif v3.1.1 algorithm (35) was used to find A/T-rich hairpin structures in the region downstream of each stop codon in medusavirus genes.

### Functional enrichment analysis

Gene ontology (GO) and KEGG Pathway enrichment analysis of the identified clusters of host genes were performed using the clusterprofiler package in R (36). KEGG pathway visualization was conducted using the pathview package in R (37).

### Data availability

The sequencing data used in this study have been submitted to the DDBJ under the accession number DRA011802.

## Results

### Transcription profile of medusavirus genes

The overall composition of the mRNA library during the course of the virus infection is shown in Fig. 1. Until 8 hpi, viral reads were less than 1% of the total reads, then they increased and reached a peak at 24 hpi. The proportion of host reads stayed at a high level during the first 8 hours, then decreased rapidly and reached a minimum at 24 hours, which still accounted for approximately half of the total reads in the library.

**Figure 1.**
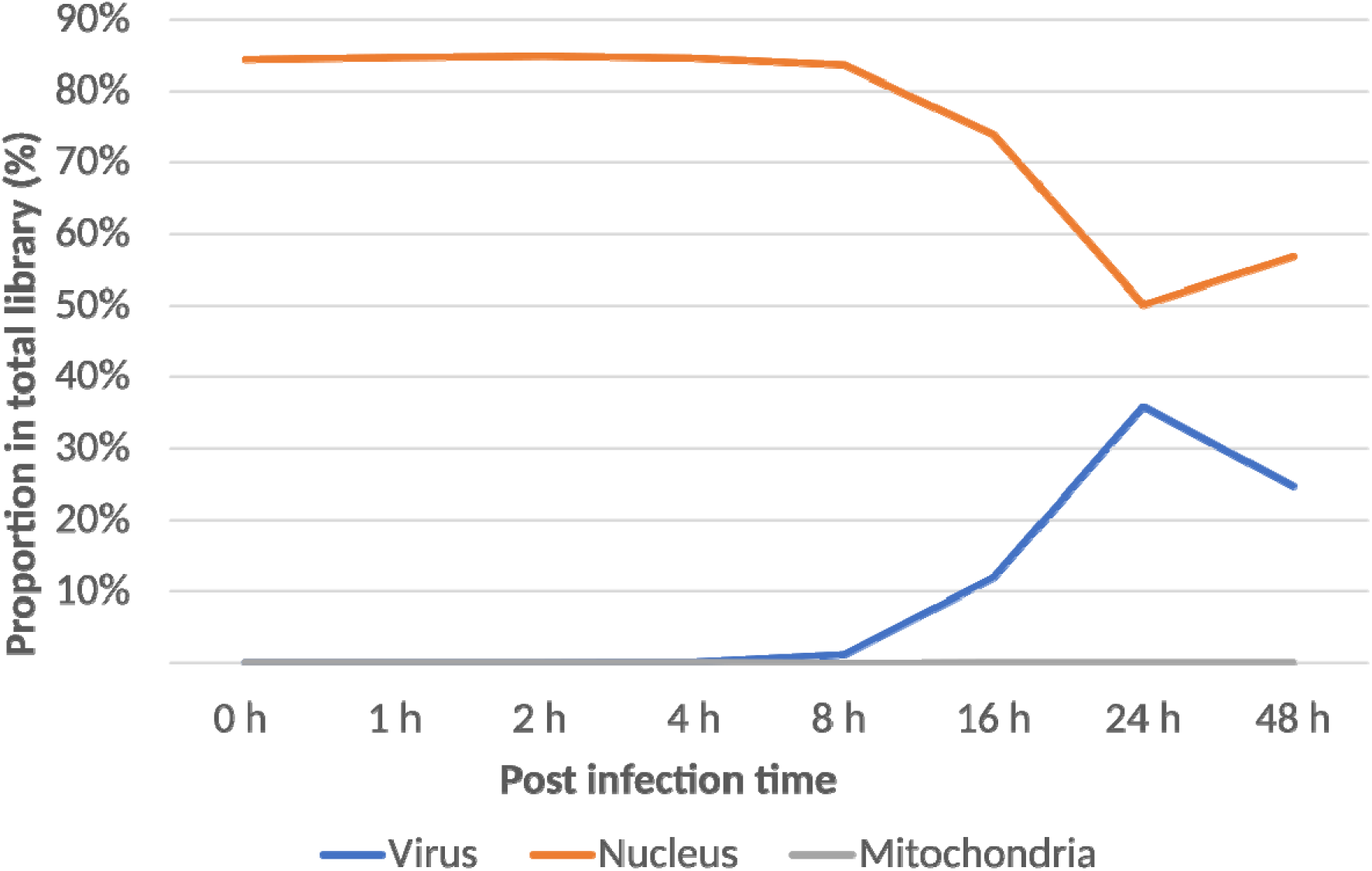
Proportions of viral and host mRNA reads at different time points during the course of the medusavirus infection.

Transcription activity was detected for all 461 viral genes (Fig. 2A). We identified five clusters of viral gene expression profiles using the k-means method (Fig. 2B) as follows: cluster 1 (“immediate early”), genes that start to be expressed immediately after the initiation of infection and before 1 hpi; cluster 2 (“early”), genes that start to be expressed at 1–2 hpi; clusters 3 and 4 (“intermediate”), genes that start to be expressed at 2–4 hpi; and cluster 5 (“late”), genes that start to be expressed at 4–16 hpi. Genes in cluster 3 showed higher z-score scaled RPKM values at 8 hpi than those in cluster 4.

**Figure 2.**
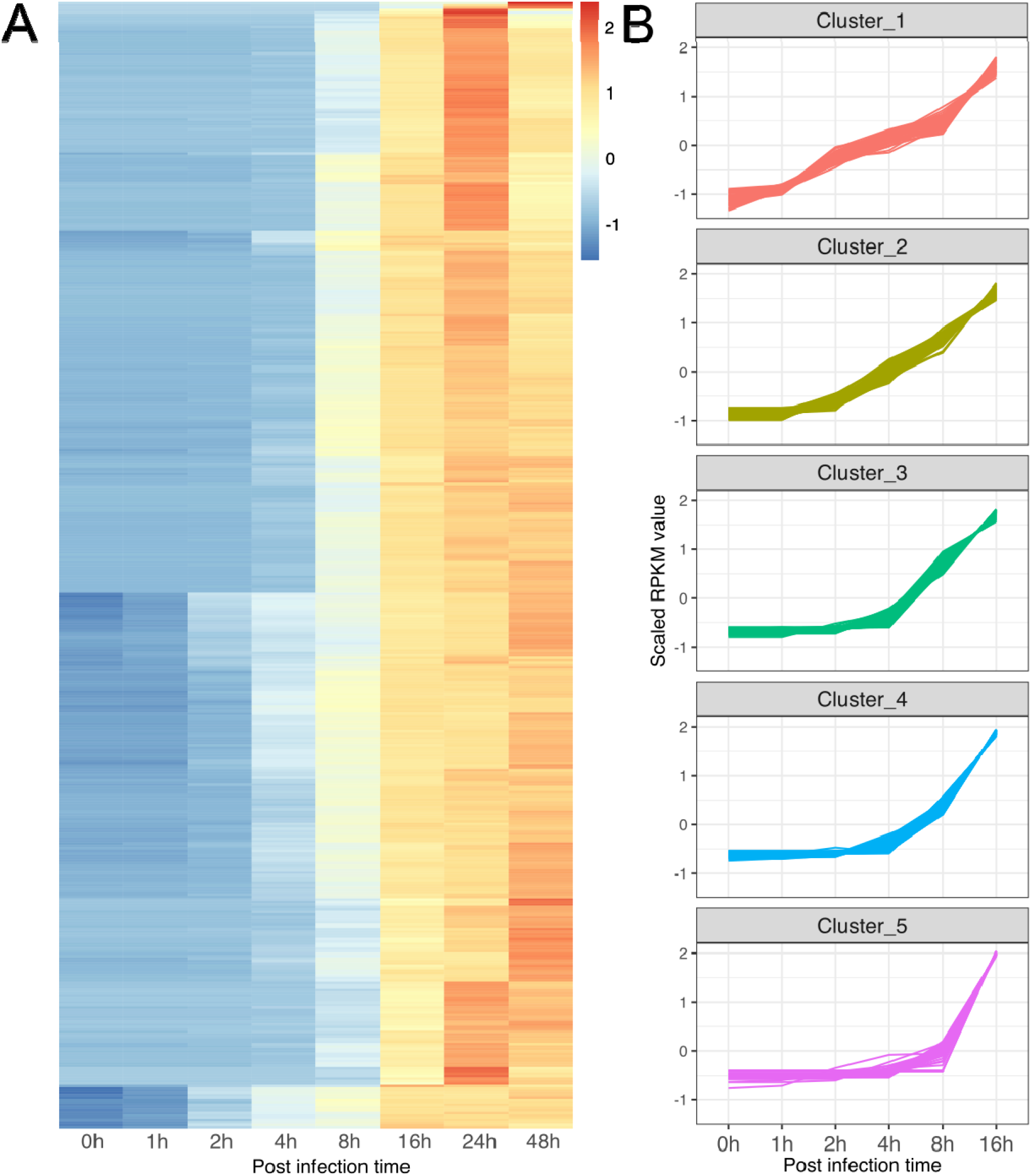
Expression of medusavirus genes at different time points during the course of the medusavirus infection. A. Heatmap of medusavirus gene expression profiles. Each column represents one time point; each row represents a viral gene; the color scale indicates z-score scaled RPKM values. B. Medusavirus temporal gene expression profiles in the five clusters. x-axis, time points post-infection; y-axis, z-score scaled RPKM value for each gene. Each line represents a viral gene.

The distribution of genes with annotated functions showed characteristic patterns among the five expression clusters (Fig. 3A–C, Supplemental Material 5). Among the 42 genes in cluster 1 (i.e., “immediate early”), 35 (83%) were unknown genes and, among the annotated genes, one was a linker histone H1 gene and another was a poly-A polymerase regulatory subunit gene. The proteins encoded by these two genes were not detected in a previous virion proteomic study of medusavirus (2). Cluster 2 included genes that were classified in the “Nucleotide metabolism” and “DNA replication, recombination, and repair” categories, including genes that encode a DNA helicase, a DNA primase, and ribonucleotide reductase large/small subunits. Cluster 3 contained genes under various functional categories, including the four core histone genes (H2A, H2B, H3, and H4), and a Yqaj viral recombinase gene and a Holliday junction resolvase gene, which may function in homologous recombination or replication. Yqaj was shown to be essential for the replication of a bacteriophage (38). Also in cluster 3 were two of five nuclease genes (“DNA replication, recombination, and repair”) as well as genes classified in “Transcription and RNA processing” (e.g., putative VLTF-2 transcription factor, putative late transcription factor 3, and transcription elongation factor S-II), “Virion structure” (e.g., major capsid protein, putative membrane protein), and “Translation" (e.g., translation initiation factor elF1 and a tRNA^His^ guanylyltransferase). Clusters 4 and 5 also contained genes under various functional categories, including those related to transcription, translation, and virion structure. Our data indicate relatively late transcription of the 79 genes that encode proteins that are known to be packaged in viral particles (Fig. 3B), and most of them (73 genes, 92%) were in the “intermediate” or “late” expression clusters (i.e., clusters 3–5).

**Figure 3.**
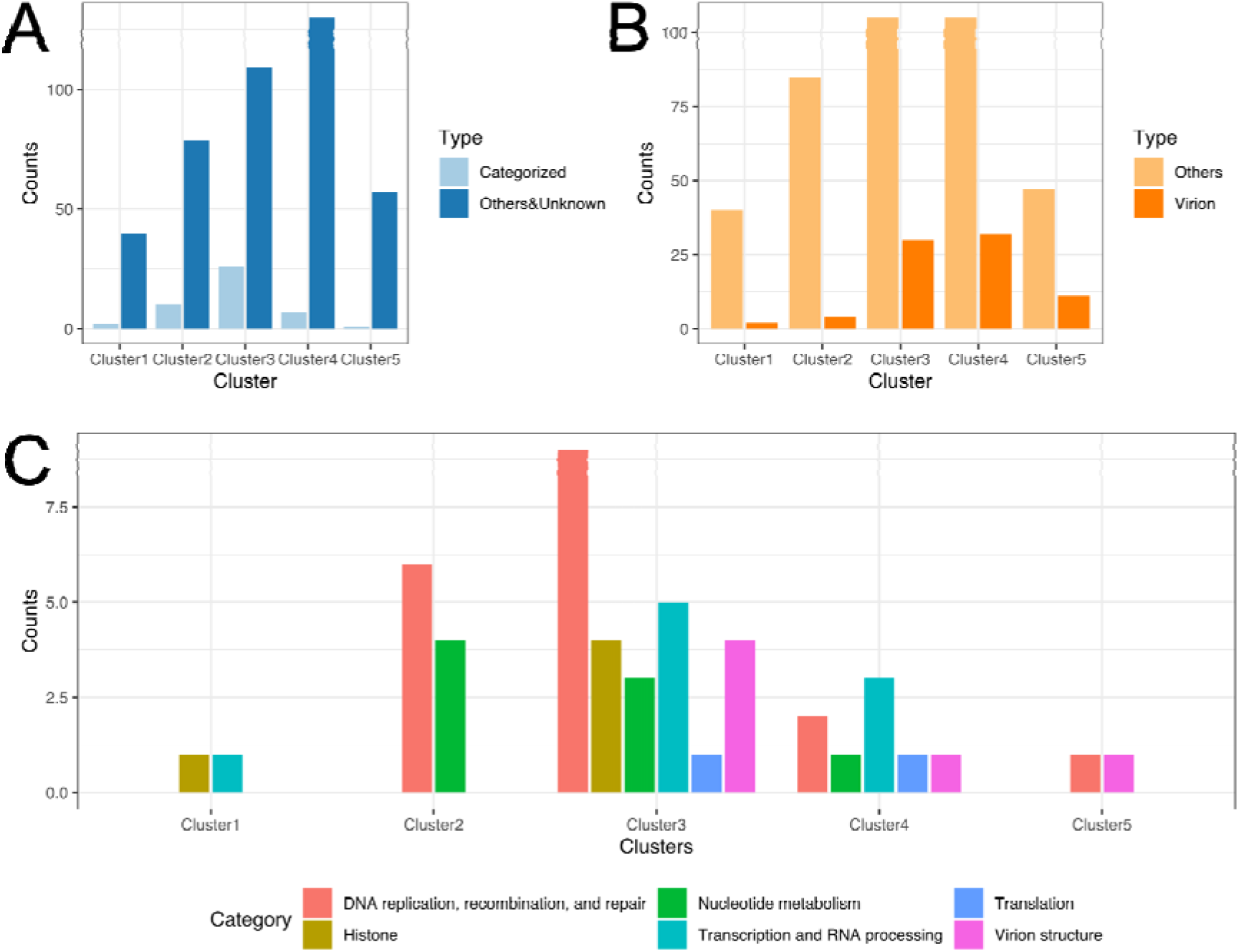
Distribution of genes with annotated functions among the five expression clusters. A. Numbers of unknown and annotated genes in the expression clusters. Light blue indicates gene with known functions; dark blue indicates genes with unknown function. B. Numbers of genes in the expression clusters. Light orange indicates genes that encode proteins not packaged in virions; dark orange indicates genes that encode proteins packaged in virions. C. Numbers of functionally annotated genes in each of the expression clusters.

### Subcellular location of viral gene products

Most of the viral gene products were predicted to target the nucleus (134 genes, 29.1%), cytoplasm (173 genes, 37.5%), mitochondrion (55 genes, 11.9%), or extracellular components (37 genes, 8%) (Fig. 4). We used the proportions of genes in these categories in each of the expression cluster to investigate the subcellular localization of their products. The proportion of nucleus-targeting proteins showed a clear descending trend, with the highest proportion in cluster 1 (45.2%) and the lowest proportions at the late stage of the infection cycle (21.2% in cluster 4; 24.1% in cluster 5). The proportion of nucleus-targeting proteins in the virion-packaged group (i.e., proteins that are known to be packaged inside the virion) was also low (21.3%). The proportion of cytoplasm-targeting proteins increased from cluster 1 (28.6%) to cluster 3 (45.9%), then decreased to cluster 5 (20.7%). Unlike the nucleus-targeting proteins, the proportion of cytoplasm-targeting proteins in the virion-packaged group was high (38.8%) and was the biggest group among the virion-packaged proteins. The proportions of mitochondrion- and extracellular-targeting proteins increased in cluster 4 (extracellular 9.5%; mitochondrion 17.5%) and cluster 5 (extracellular 13.8%; mitochondrion 19.0%), however, their proportions in the virion-packaged group was low (5 genes, 6.3% of the total virion proteins). Smaller numbers of proteins were predicted to target cell membrane (23 genes), endoplasmic reticulum (16 genes), peroxisome (13 genes), Golgi apparatus (6 genes), and lysosome/vacuole (4 genes). Their proportions and absolute numbers increased in the “intermediate” or “late” expression clusters (i.e., clusters 3–5) (Supplemental Material 3). In addition, approximately half of the peroxisome- (6 out of 13 genes) and cell membrane-targeting (11 out of 23 genes) proteins were in the virion-packaged groups.

**Figure 4.**
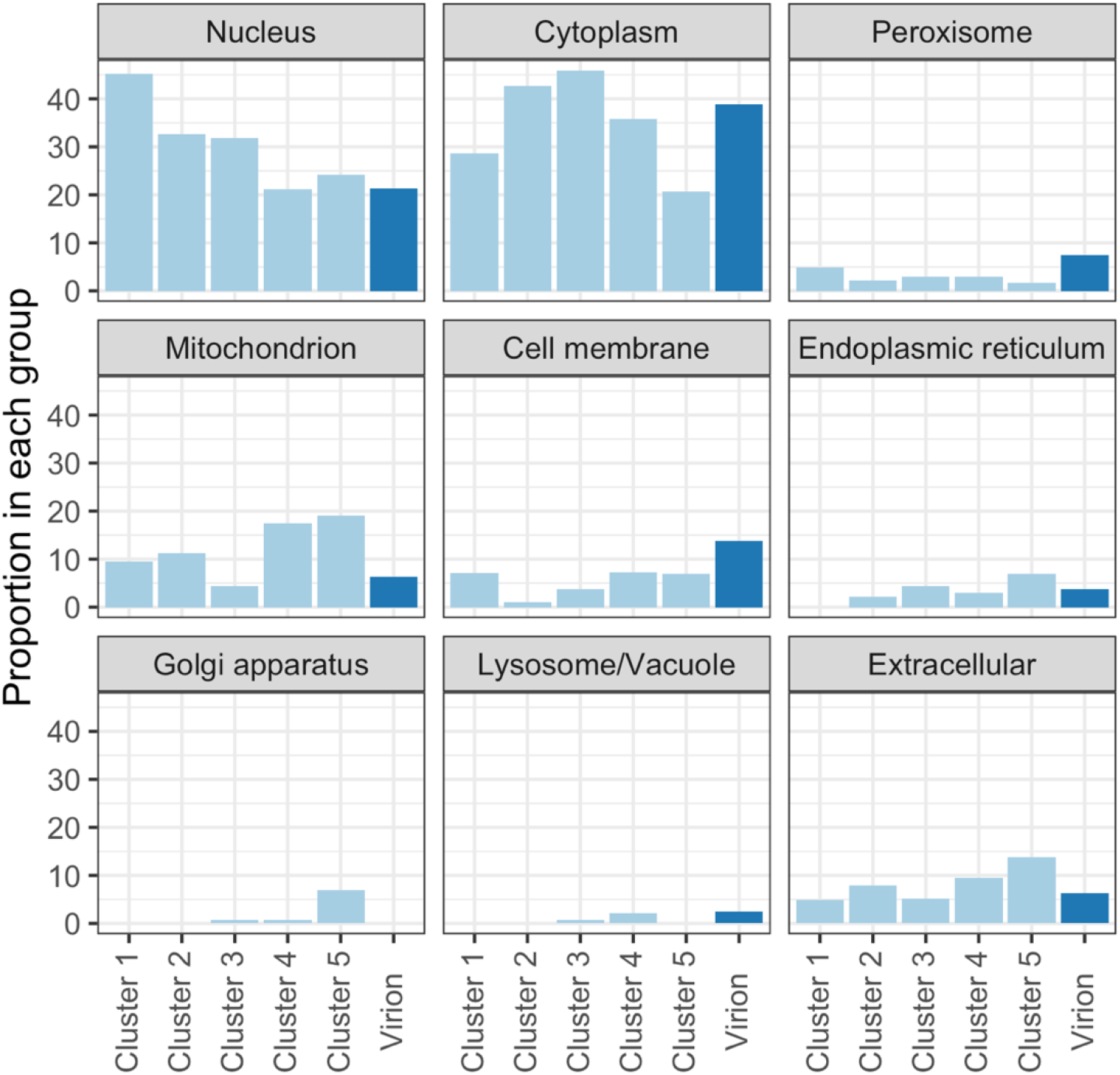
Predicted subcellular locations of the products of viral genes. The height of each bar indicates the proportion of genes in each cluster. Light blue indicates the proportion of genes in the expression clusters; dark blue indicates the proportion of genes among the viral genes whose products are known to be packaged inside the virion.

### Putative regulatory elements

To investigate the regulatory mechanisms of medusavirus gene expression, we analyzed the genomic localizations of the temporal gene expression clusters and associated gene functions. However, this analysis did not detect any definitive features related to the organization of genes in the genome and their temporal or functional groups (Fig. 5). We then performed *de novo* motif searches in the 5′ region upstream of the viral genes and identified five motifs that were statistically significantly overrepresented (*p* <1×10^−5^, Supplemental Material 4), namely GCCATRTGAVKTCATRTGGYSRSG (palindromic motif, 53 occurrences), VMAAMAAMARMAAMA (poly-A motif, 251 occurrences), GCCRYCGYCGH (GC-rich motif, 134 occurrences), NRAAWAAA (AATAAA-like motif, 123 occurrences), and GTGTKKGTGGTGGTG (GT-rich motif, 37 occurrences) (Fig. 6; Supplementary Material 4, Table 1&2).

**Figure 5.**
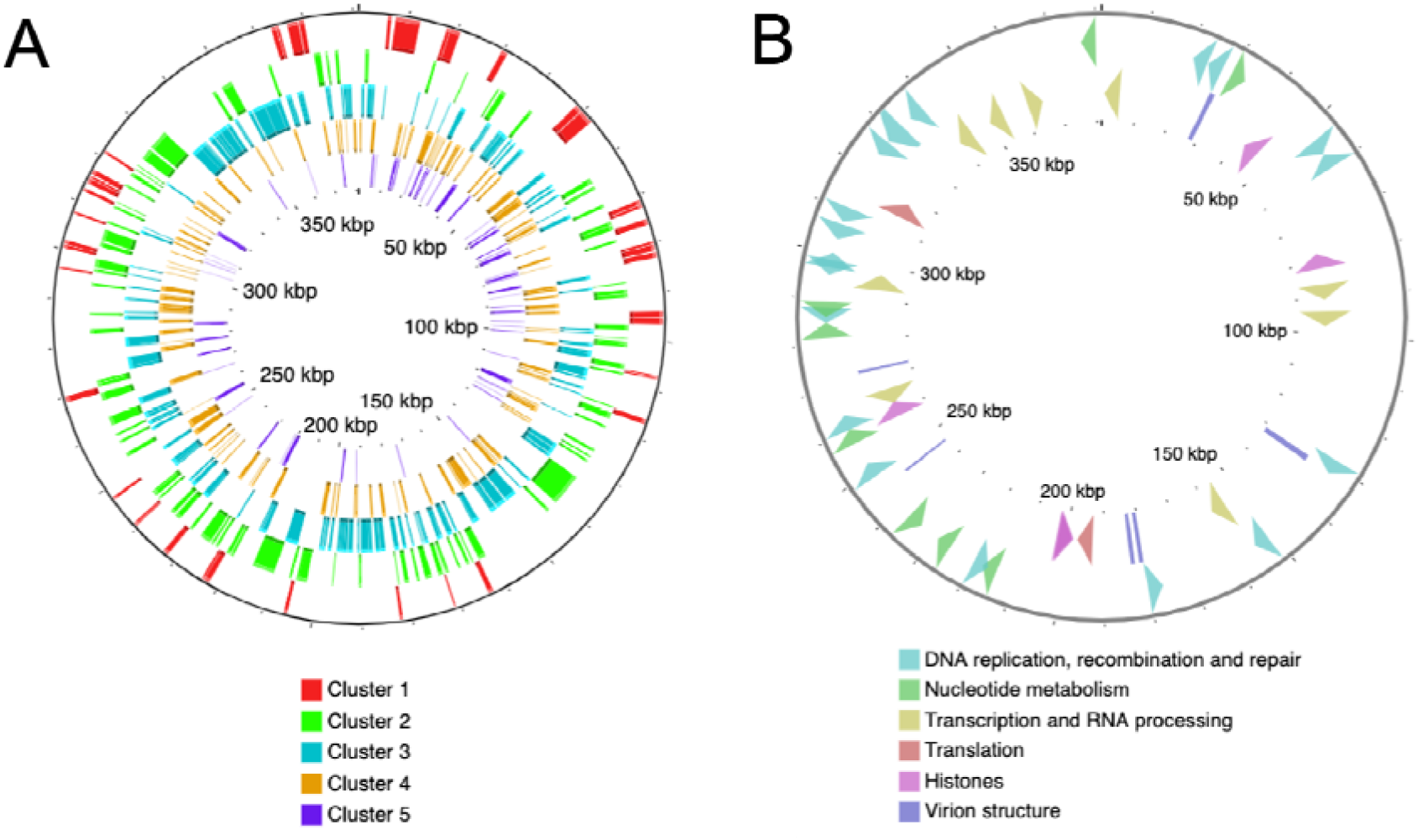
Organization of genes in the medusavirus genome. A. Organization of the five expression clusters on the viral genome. B. Organization of functional groups of genes on the viral genome. Outside layer: genes classified in the “DNA replication, recombination, and repair” and “Nucleotide metabolism” categories; Inside layer: Genes classified in the “Transcription and RNA processing”, “Translation”, “Histones”, and “Virion structure” categories.

**Figure 6.**
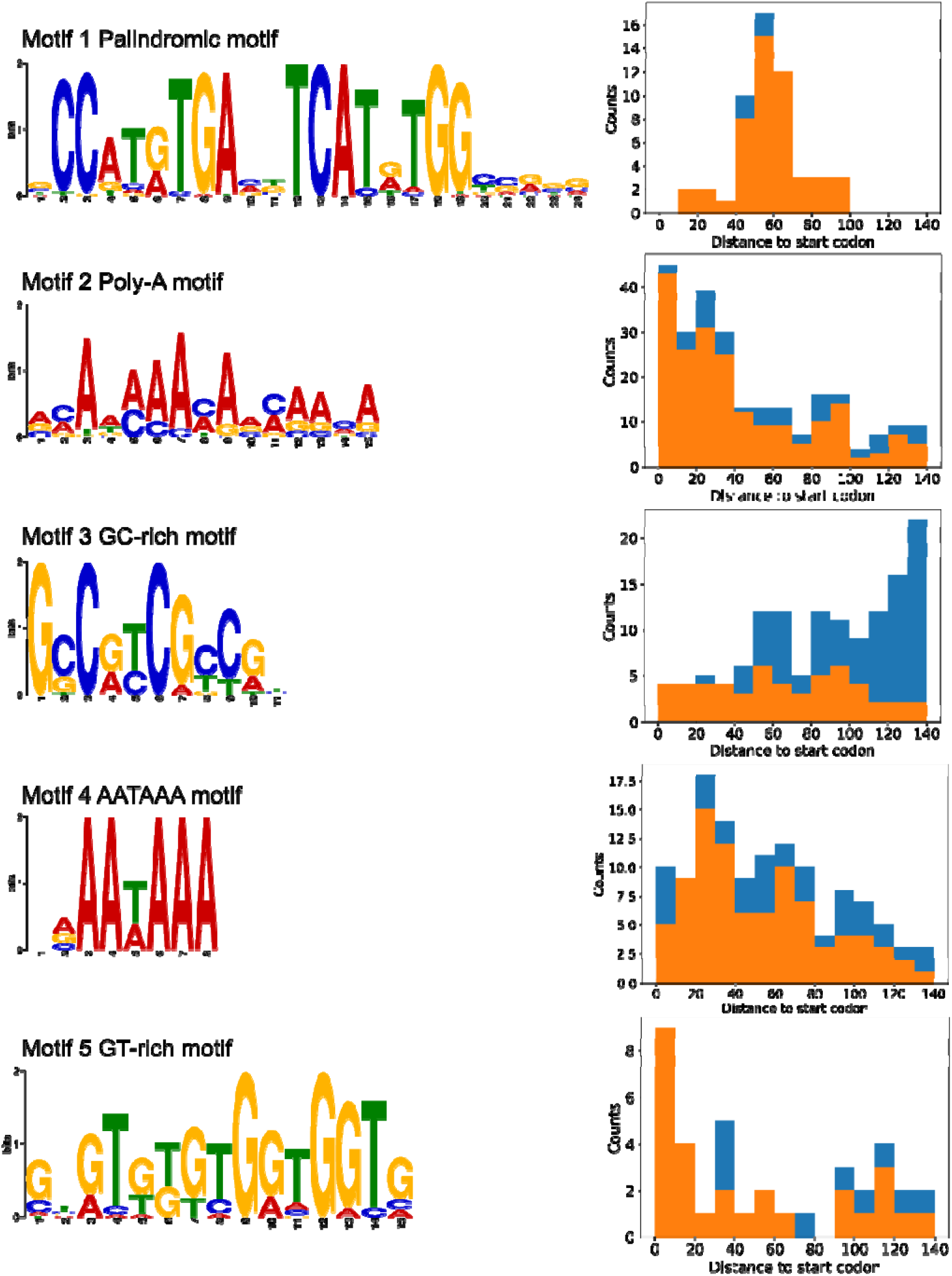
Sequence motifs enriched in the 5′ region upstream of the genes in the medusavirus genome and their distribution relative to the corresponding start codon. Left panel, motif name and its LOGO; right panel, distance to corresponding start codon; orange indicates motifs that did not overlap with neighboring genes; blue indicates motifs that overlapped with neighboring genes.

**Table 1.** Distribution of upstream motifs in intergenic regions^*a*^ of the medusavirus genome.

|  | Motif 1 | Motif 2 | Motif 3 | Motif 4 | Motif 5 |
| --- | --- | --- | --- | --- | --- |
| Total | 48 | 152 | 807 | 19 | 253 |
| IR<sup>b</sup> | 30 | 84 | 43 | 19 | 27 |
| Background frequency<sup>c</sup> | 0.105 |  |  |  |  |
| *p*-value | 4.60e-18 | 5.62e-42 | 1.00 | 0.132 | 0.495 |

<sup>a</sup>A binomial test was used to assess whether each motif was preferentially located in intergenic regions (IRs).
<sup>b</sup>Only motifs that did not overlap with predicted genes were considered to be located in IRs.
<sup>c</sup>Background frequency was calculated by dividing the length of all the IRs by the length of the whole genome.

**Table 2.** Preference of the palindromic and poly-A motifs for up- or down-stream regions of the medusavirus genes^*a*^

|  | Motif 1 Palindromic |  | Motif 2 Poly-A |  |
| --- | --- | --- | --- | --- |
|  | With Motif | Without Motif | With Motif | Without Motif |
| Divergent<sup>b</sup> | 22 | 93 | 40 | 75 |
| Convergent<sup>c</sup> | 0 | 88 | 4 | 84 |
| *p*-value | 2.76e-06 |  | 8.48e-08 |  |

<sup>a</sup>Only motifs predicted to be located in intergenic regions were used to determine their preferred location. The *p*-values were calculated by the Fisher exact test.
<sup>b</sup>Divergent cases were defined as motifs being located in the upstream regions of both neighbor genes.
<sup>c</sup>The convergent cases were defined as the motifs being located in the downstream region of both neighbor genes.

The palindromic motif was preferentially found in the region approximately 40–70 bp upstream of the start codon. The poly-A motif was preferentially found in the region approximately 0–40 bp upstream of the start codon. The GC-rich motif had no obvious preferred position upstream of the start codon, but it often overlapped with upstream genes. The AATAAA motif was preferentially found in the region approximately 0–60 bp upstream of the start codon, which is similar to the preferred positions of the poly-A motif. The GT-rich motif was preferentially found close to the start codon (Fig. 6).

The palindromic motif was highly associated with the genes in cluster 1 (“immediate early”) (Fig. 6). Among the 53 viral genes with the upstream palindromic motif, 40 (75%) were in cluster 1, and they made up 95% of the genes in this cluster. The 13 other genes with the palindromic motif were distributed among the other clusters; 8 in cluster 2 and the other 5 in clusters 3, 4, and 5 (Fig. 7). Furthermore, among the 56 genes detected at 1 hpi, 49 (88%) had the palindromic motif. We also found that 81.0% of the genes in cluster 1 and 69.7% of the genes in cluster 2 had the upstream poly-A motif, whereas they made up only 40.2%–55.2% of the genes in clusters 3– 5 (Fig. 7). For the other motifs, we found no specific association with gene clusters.

**Figure 7.**
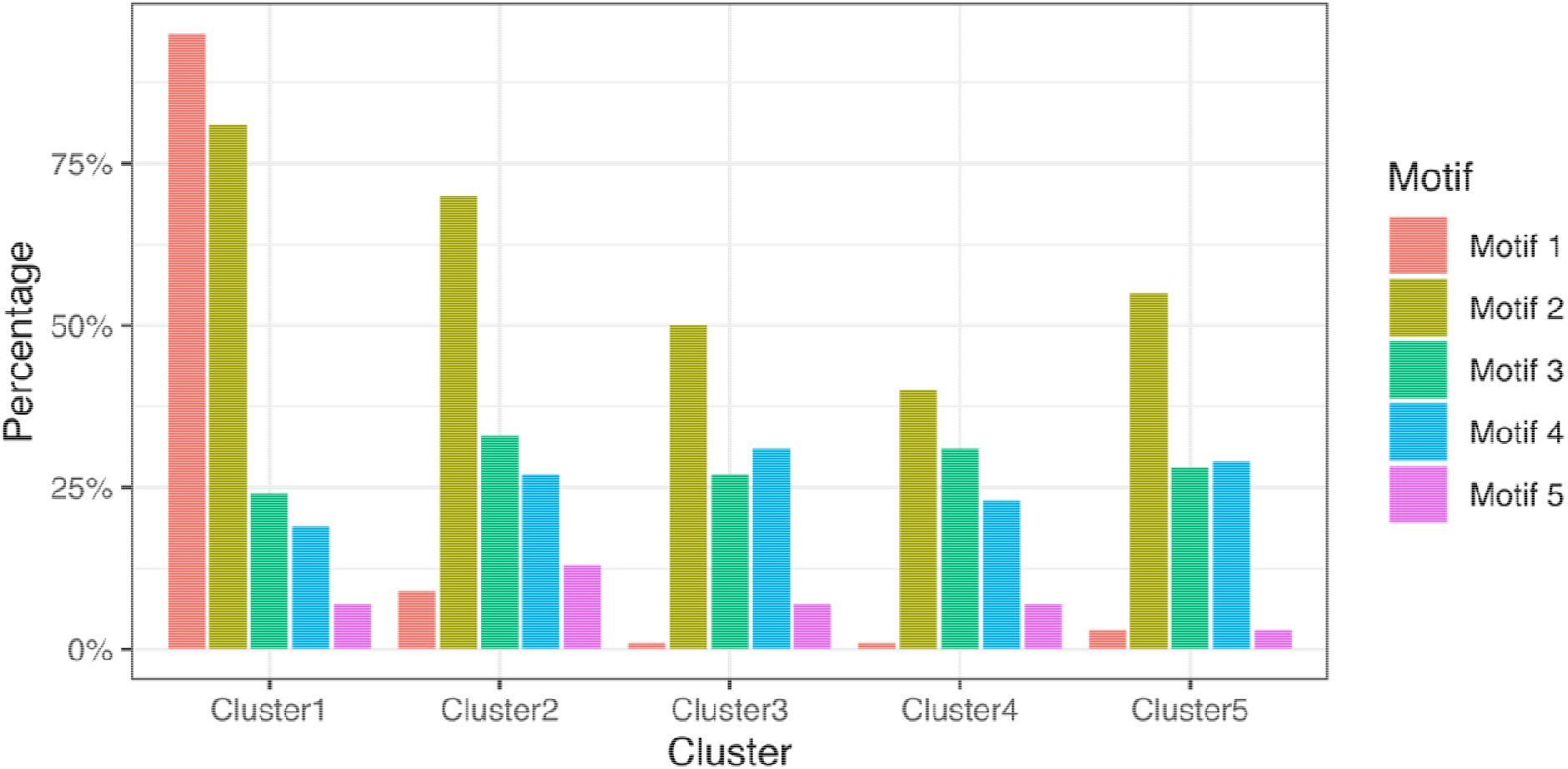
Proportion of genes with the different upstream motifs in each expression cluster.

To investigate if these upstream motifs were promoter motifs, we scanned the medusavirus genome with these motifs. The palindromic and poly-A motifs were statistically significantly more abundant in intergenic regions than in coding sequences (*p* <2.2×10^−16^, Table 1). Furthermore, these two motifs were more frequent in the upstream intergenic regions of genes than in the downstream intergenic regions (*p* <10^−5^, Table 2). The other three motifs showed no preference for either intergenic regions or coding sequences. We also searched the 3′ downstream regions of the medusavirus genes for hairpin structures but failed to identify any. This is unconsistent to previous reports of hairpin structures found in *Acanthamoeba polyphaga mimivirus* (stem length ≥13 bp, loop ≤5 bp), *Megavirus chilensis* (stem length ≥15 bp), *Pithovirus sibericum* (stem length ≥10 bp, loop ≤10 bp), and virophage sputnik, noumeavirus, and melbournevirus in the family *Marseilleviridae* (3, 21, 39–41).

### Host nuclear transcriptional profile was greatly altered

The proportion of host mRNA reads and their expression levels assessed by RPKM did not show large changes until 8 hpi (Figs. 1 and 8A). After 8 hpi, the proportion of host reads decreased rapidly and the proportion of viral reads increased. Our cluster analysis of the dataset of 0–16 hpi showed that the transcription profile of the host *A. castellanii* genes changed greatly between 8 hpi and 16 hpi, with two expression clusters for the host genes (Fig. 9A). Of the 10,627 *A. castellanii* genes examined, 7970 (75%) were in cluster 1. Their expression levels decreased across time, especially between 8 hpi and 16 hpi. The remaining 2657 (25%) genes were in cluster 2, and their expression levels increased (at 16 hpi, mean log2 fold-change are −0.445 and 0.364 for cluster 1 and cluster 2, respectively).

**Figure 8.**
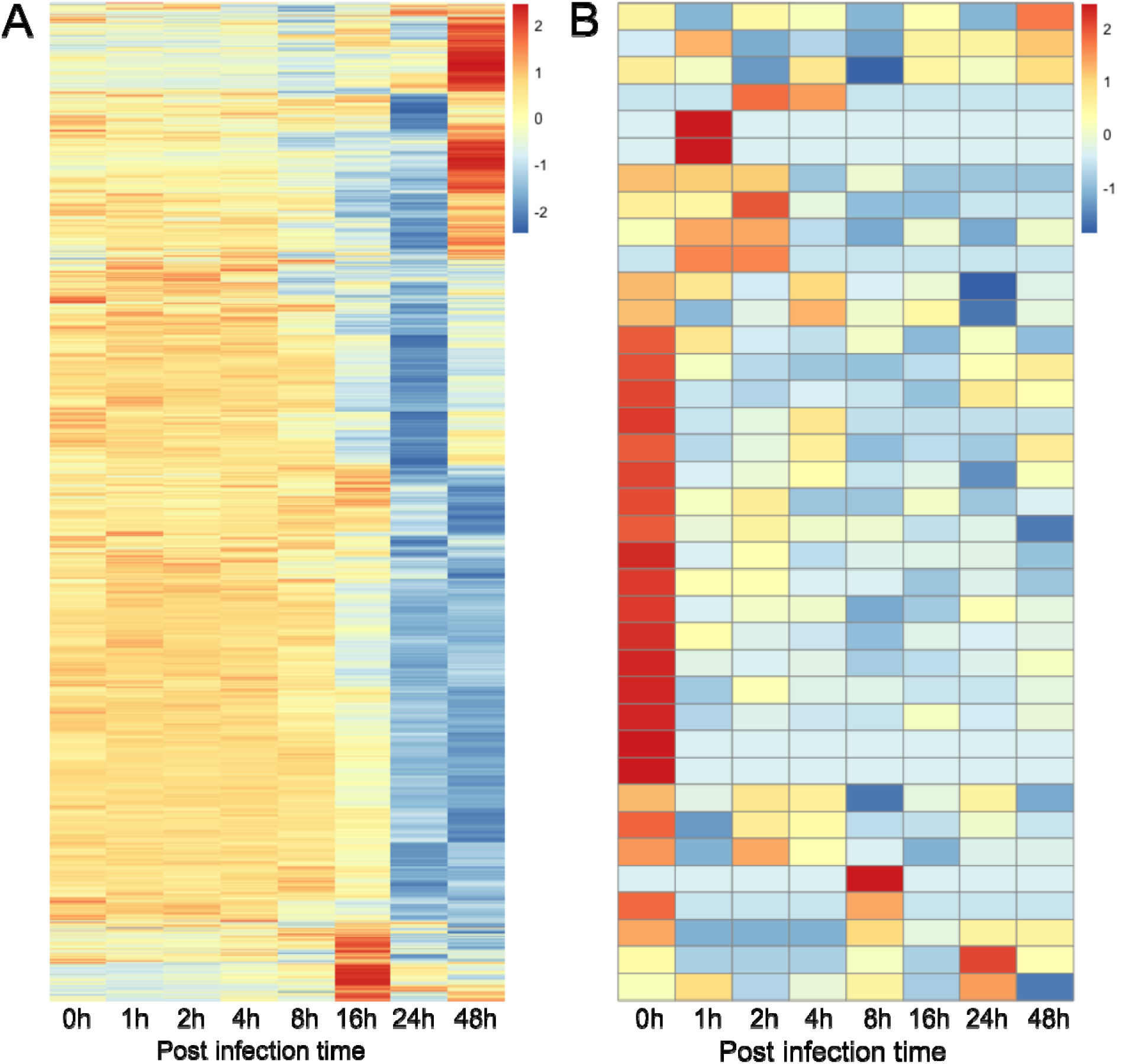
Transcription profiles of the *Acanthamoeba castellanii* (host) genes. A. Host nuclear genes. B. Host mitochondrial genes. X-axis, time points of the infection cycle; Y-axis, different genes in the host genome and its mitochondrial genome. The color scale indicates z-score scaled RPKM values.

**Figure 9.**
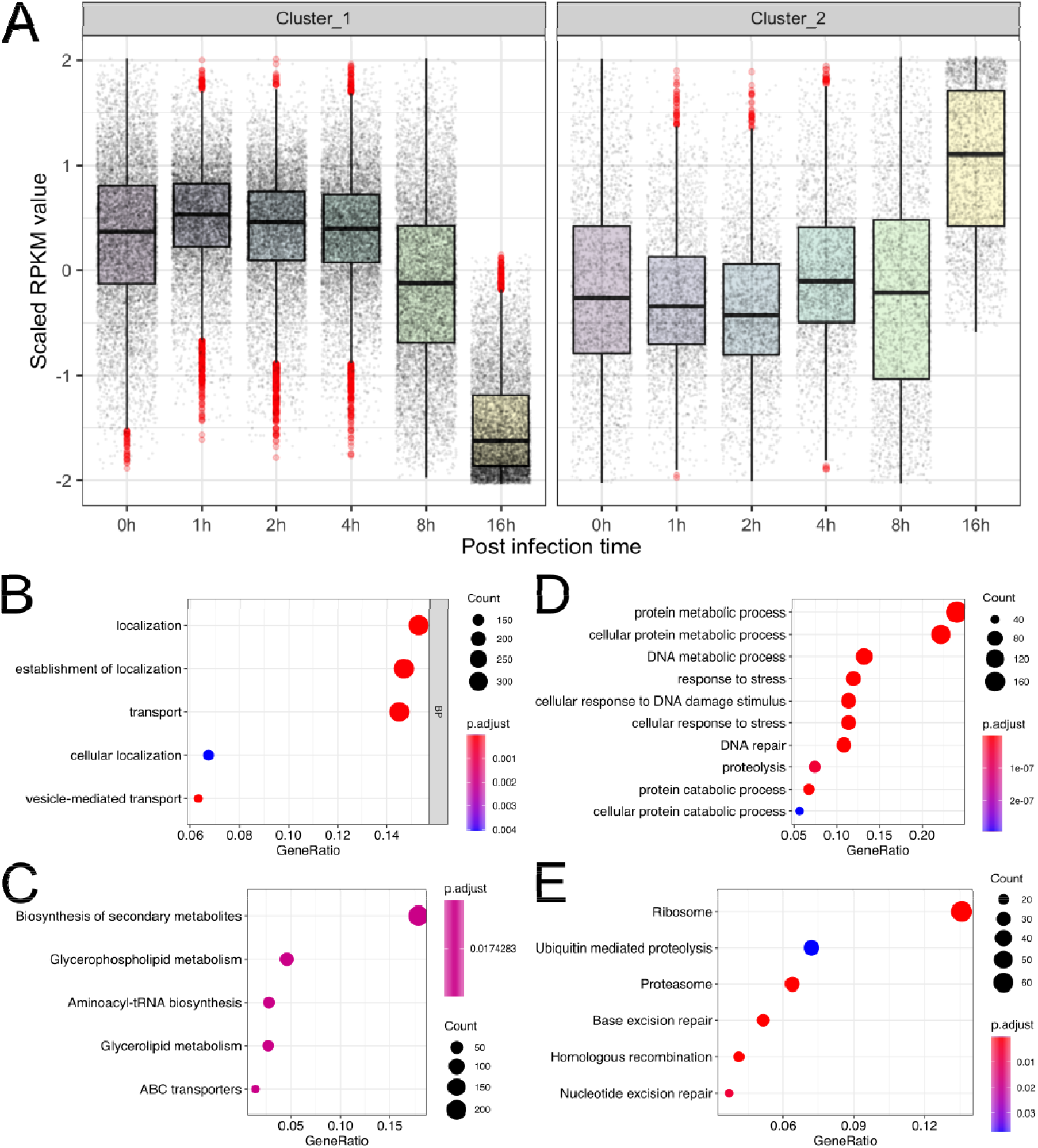
*Acanthamoeba castellanii* (host) nuclear gene expression clusters and their predicted functions. A. Two expression clusters were identified for the host nuclear genes. B. Enriched GO terms for the genes in cluster 1. C. Enriched KEGG pathways for the genes in cluster 1; D. Enriched GO terms for the genes in cluster 2. E. Enriched KEGG pathways for the genes in cluster 2.

Based on the host gene clusters, we performed GO and KEGG Pathway functional enrichment analyses (Fig. 9B–E). Cluster 1 genes were enriched in cellular transportation-related GO terms, such as “localization”, “establishment of localization”, and “transport” (Fig. 9B). Cluster 2 genes were enriched in 60 different GO terms that fell into two main categories (Fig. 9D, Supplemental Material 6). One category comprised terms related to “cellular protein metabolic process” and “proteolysis involved in cellular protein catabolic process”, and the other category comprised stress-related terms such as “DNA repair”.

Cluster 1 host genes were also enriched in KEGG pathways, including “biosynthesis of secondary metabolites”, “glycerophospholipid metabolism”, “aminoacyl-tRNA biosynthesis”, “glycerolipid metabolism” and “ABC transporters” (Fig. 9C). Cluster 2 host genes were enriched with “Ribosome”, “Ubiquitin mediated proteolysis”, “Proteasome”, “Base excision repair”, “Homologous recombination”, and “Nucleotide excision repair” (Fig. 9D). Clearly, the first three of these pathways correspond to the enriched GO terms in cluster 2 related to protein catabolic and metabolic processes, and the latter three pathways correspond to the enriched GO terms related to DNA repair and stress response.

### Host nuclear genes expression pattern changed after 16 hpi

Clusters identified in the first 16 hpi did not maintain their expression pattern after 16 hpi (Supplemental Material 6, Fig. 2). The expression levels of some genes annotated with GO term “transport” were increased greatly at 48 hpi, indicating that proteins encoded by genes involved in membrane-related transport, such as the v-SNARE protein family, Sec23/24 trunk domain-containing protein, and Sec13, were highly abundant in the late stages of infection (Supplemental Material 6, Fig. 3). In contrast, cluster 2 genes, which were activated at 16 hpi, were suppressed at 24 hpi, then recovered to some extent at 48 hpi. We found that some of the genes that were activated at 48 hpi were encystment-mediating genes, which included an encystation-mediating serine proteinase (EMSP), eight cysteine protease proteins, cyst specific protein 21 (CSP21), and two cellulose synthases (42–46) (Supplemental Material 6, Fig. 4).

### Mitochondrial expression was maintained during medusavirus infection

In our RNA-seq data, 37 of the 53 genes encoded in the *A. castellanii* mitochondrial genome had at least one read count during the course of the infection (Fig. 8B). The numbers of mitochondrial mRNA reads were low (4365 reads at 0 hpi and maintained <2500 at the later time points). However, the numbers of mitochondrial reads were more stable than the numbers of reads that were mapped onto the nuclear genome, the latter of which decreased by about 30% during the course of the infection.

Mitochondrial genes with the highest read counts were related to ribosomal RNA genes (AccaoMp41, 23S-like ribosomal RNA and AccaoMp42, 16S-like ribosomal RNA), followed by energy metabolism (AccaoMp13, H (+)-transporting ATPase subunit 9). The proportions of the other mapped mitochondrial mRNA reads were small. Among the 16 mitochondrial genes that were not detected in the RNA-seq data, 11 were tRNA genes, 3 were ribosomal protein genes, and 2 genes were of unknown function.

## Discussion

We performed RNA-seq to dissect the transcriptional program of the medusavirus. Medusavirus has been reported to initiate its genome replication in the host nucleus and maintain the nuclear membrane intact during the infection cycle, with occasional induction of the encystment of the host amoeba *A. castellanii* at approximately 48 hpi (2). We found that transcription began in less than one hour after the start of infection even though medusavirus has no RNA polymerase genes. Compared with other amoeba-infecting viruses, the speed of the medusavirus infection cycle was slow and apparently weak because it took approximately 24 hours for the virus to reach its expression peak, which was still only about 35% of the total mRNAs (MOI = 2.88). In contrast, mimivirus and marseillevirus genes occupy 80% of the total mRNA library at less than 6 hpi (MOI = 100 and 1000 for marseillevirus and mimivirus, respectively) (24, 25). The slow and mild medusavirus infection may be explained by the different MOI used in the infection experiments, where higher MOI has been reported to accelerate the infection course (47, 48). An additional explanation may be a slow start of medusavirus replication. While mimivirus carries its DNA and RNA polymerases in the viral particles, which is considered to help in forming replication centers as soon as the viral particles open up (49), medusavirus not only does not encode many transcription- and translation-related genes, but also does not carry DNA polymerase in viral particles. Thus this essential enzyme needs to be synthesized after infection, which may account for its slow infection.

Clustering of the medusavirus gene expression profiles showed clear temporal expression patterns akin to those observed for other giant viruses (24, 25, 50). We found that linker histone H1 was transcribed immediately after the beginning of transcription. This histone may cooperate with high-mobility group proteins in viral particles to regulate the accessibility of the viral genome for the subsequent transcription process (51, 52), or it may function to regulate the host chromatin. Notably, the viral linker histone H1 gene showed a different expression pattern from the genes for the four core histones, which were expressed later and packaged into viral particles. These results suggest different functional roles between the linker histone H1 and the core histones.

The predicted subcellular localization of the viral gene products showed that cluster 1 had a higher proportion of nucleus-targeting genes than the other clusters. The proportions of nucleus-targeting genes in the medusavirus and medusavirus stheno genomes were ranked 6^th^ and 4^th^, respectively, among all known amoeba-infecting NCLDVs, which indicated the importance of remodeling the nuclear environment immediately after medusavirus infection (Supplemental Material 4, Fig. 1). The remodeling probably contributes to subsequent viral gene transcription and DNA replication within the host nucleus. Putative cytoplasm-targeting genes were enriched in the virion-packaged group (31 genes, 38.8% of genes in virions) and almost half of the cell membrane and peroxisome-targeting genes were also packaged inside virions, suggesting that there may be interactions between virion-packaged proteins and the host cytoplasm at an early phase of infection. The increasing expression of genes targeting mitochondrion, endoplasmic reticulum, and Golgi apparatus suggests that the medusavirus synthesizes these genes, probably to maintain or reprogram the functions of these organelles, after starting infection rather than bringing them with the virions.

The enrichment of the palindromic motif in the upstream region of genes that were transcribed immediately after infection suggests that this motif may be the immediate early promoter of medusavirus genes. The poly-A motif that we detected in the upstream region of early expressed genes is reminiscent of the A/T-rich early promoter motifs found in other giant viruses in the phylum *Nucleocytoviricota* that have been proposed to have a common ancestral promotor motif, TATATAAAATTGA (53–56). The poly-A motif in the upstream regions of the medusavirus genes also may have evolved from this common ancestral motif. Although the AATAAA motif was not preferentially located in the intergenic regions of the medusavirus genome, it is similar to the 3′-end motif in the polyadenylation signal sequence in eukaryotes (57). The AATAAA motif also was detected in mimivirus but it did not function as a polyadenylation signal (39). Regarding the 3′-end processing mechanism of giant viruses, A/T-rich hairpin structures have been detected after stop codons (3, 21, 39–41) and proteins that can recognizing these structure have been studied (58). However, we did not find any A/T-rich hairpin structures in the 3′ downstream regions of medusavirus genes.

We identified two temporal clusters for host genes during viral replication. The decrease of RPKM of most (75%) host genes at 16 hpi suggests that the host genes experienced global suppression. GO terms related to “localization” and “transport” were enriched in host cluster 1, suggesting that decreased transport activity occurred within the host cell during the course of the virus infection. In addition, the increased representation of KEGG pathways “Ribosome” and “Proteosome” and GO terms “cellular protein metabolic process” and “proteolysis involved in cellular protein catabolic process” suggested increased ribosome and proteolysis activity during the course of the virus infection.

The GO terms “DNA repair” and “cellular response to stress”, which correspond to KEGG pathways “Homologous recombination”, “Base excision repair”, and “Nucleotide Excision Repair”, were enriched at 16 hpi (Fig. 9C, D). The genes with these annotations may be used by the host *A. castellanii* to recognize DNA damage. However, because new virions are detected at approximately 14 hpi, 16 hpi may be the end of the first-round of viral infection and the beginning of the second round (2). Therefore, the activation of these genes may be related to the early events in viral infection. The ataxia-telangiectasia-mutated (ATM) pathway is a key pathway that recognizes DNA damage and induces homologous recombination repair. In the ATM pathway, the Mre11-Rad50-Nbs1 complex detects damaged DNA by interacting with double-strand and single-strand breaks. This results in the activation of ATM and CtIP, which promotes the resection of double-strand breaks (59, 60). Then, downstream genes in the pathway promote filament formation (replication factor A2 (RPA), and factors Rad51 and Rad52), strand invasion (Rad54), and DNA synthesis (DNA polymerase δ subunit 1) (60–64). The expression levels of most of the genes in the ATM pathway were increased at 16 hpi (Fig. 10). Activation of the ATM and ATR signaling pathways and homologous recombination repair have been reported to aid polyomavirus reproduction (simian virus 40 and JC polyomavirus) (60, 65, 66). Medusavirus may also make use of cellular homologous recombination apparatuses for genome replication.

**Figure 10.**
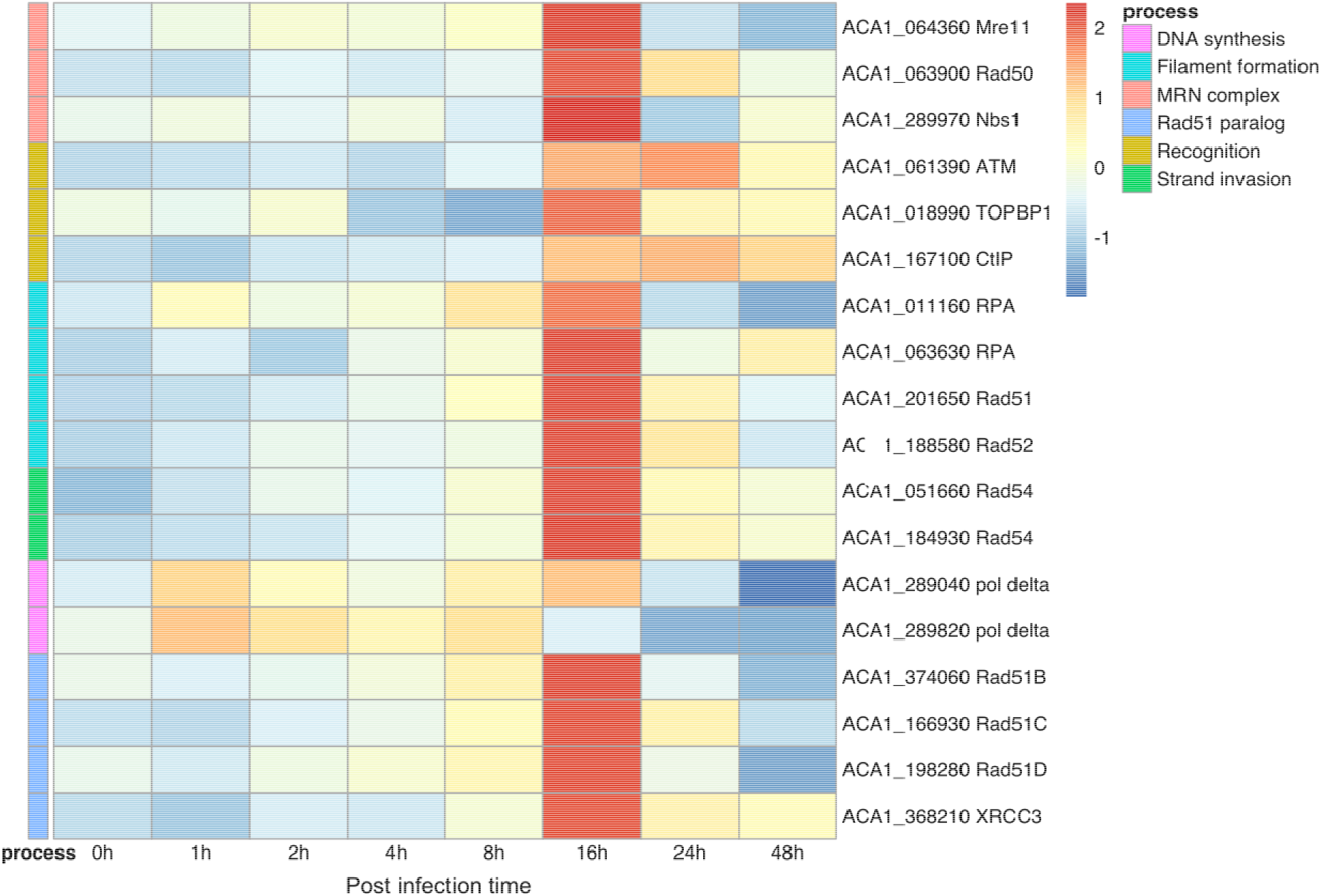
Expression profile of genes related to homologous recombination repair (KEGG pathway accession number: acan03440). The color scale indicates z-score scaled RPKM values. The process bar on the left indicates the process associated with each gene. Gene names and locus tags are on the right side.

The trend in the host gene expression patterns during the first 16 hpi were not maintained until 48 hpi and the viral reads decreased from 24 to 48 hpi. Some of the *A. castellanii* cells forming cyst-like structures may account for this change. The encystment of *A. castellanii* has been proposed as an effective way for it to survive in undesirable growth conditions such as the presence of toxic chemicals, shortage of nutrients, or virus infection (67–69). Given the presence of activated encystment-related genes at 48 hpi (Supplemental Material 6, Fig. 4), it is tempting to speculate that encystment helps *A. castellanii* defend itself against medusavirus infection.

The expression pattern of the *A. castellanii* mitochondrial genes during the course of the medusavirus infection was similar to the pattern found in marseillevirus (25). All tRNA-encoding genes had low expression levels, possibly because transcripts with a poly-A tail were used to build the RNA library and tRNA genes do not have poly-A tails. The genes with the highest expression levels included genes involved in energy metabolism and ribosomal RNA genes. However, unlike the read counts for the host nuclear genes, the transcriptional activity of the mitochondrial genes was maintained, suggesting a stable energy supply was available for virus infection. In conclusion, the expression levels of the mitochondrial genes were maintained during the course of the medusavirus infection and genes related to ribosomal RNA and energy metabolism have a relatively high value.

In summary, our transcriptome data were clearly delineated into five temporal expression clusters for viral genes. Most of the immediate early genes (cluster 1) were uncharacterized genes and had a palindromic promoter-like motif upstream of their start codons. Many of the immediate early gene products were predicted to target the host nucleus, suggesting that the medusavirus modifies the host nuclear environment after the start of infection, and involves the action of dozens of genes. The genes that were expressed later (clusters 2–5) have various functions. The viral histone H1 gene is in the cluster 1, whereas the four core histone genes are in cluster 3, suggesting they have distinct roles in viral replication. The transcriptional landscape of the host nuclear genes was altered during infection especially after 8 hpi. At 16 hpi, the host nuclear transcriptional showed a great alteration. Our transcriptome data will serve as a fundamental resource for further characterizing the infection strategies of medusaviruses, which are a group of amoeba-infecting giant viruses that have no close relatives among the diverse NCLDVs.

## Acknowledgments

Computational time was provided by the SuperComputer System, Institute for Chemical Research, Kyoto University. This work was supported by JSPS/KAKENHI (Nos. 18H02279 and 20H03078), The Kyoto University Foundation, and the Collaborative Research Program of the Institute for Chemical Research, Kyoto University (Nos. 2018-32, 2019-34, and 2020-31). We thank Margaret Biswas, PhD, from Edanz Group (https://en-author-services.edanz.com/ac) for editing a draft of this manuscript.

