## Supplementary Material 4 for "RNA-seq of the medusavirus suggests remodeling of the host nuclear environment at an early infection stage"

**Supplementary File 5**

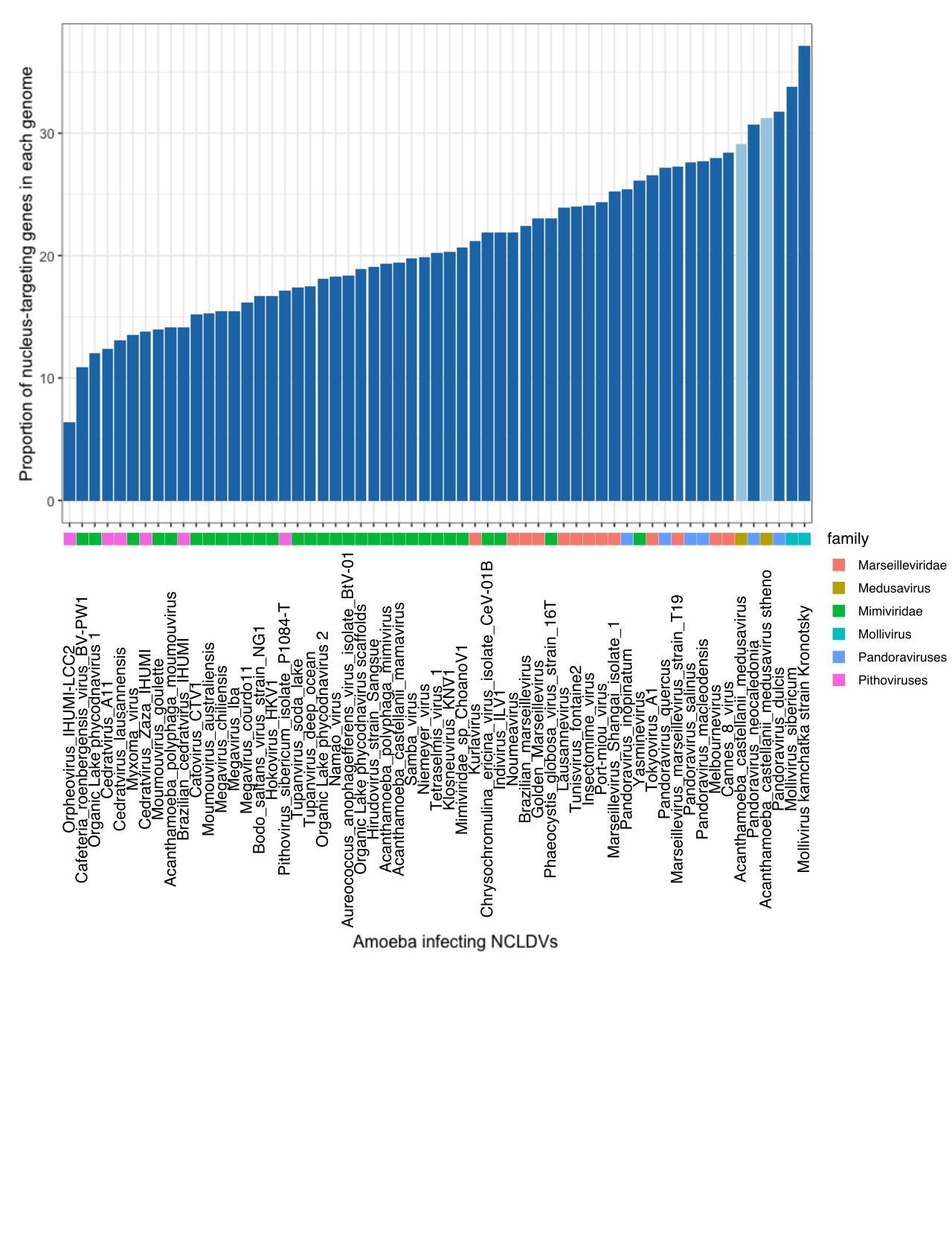

Fig.1 Deeploc prediction results of other amoeba infecting NCLDVs.

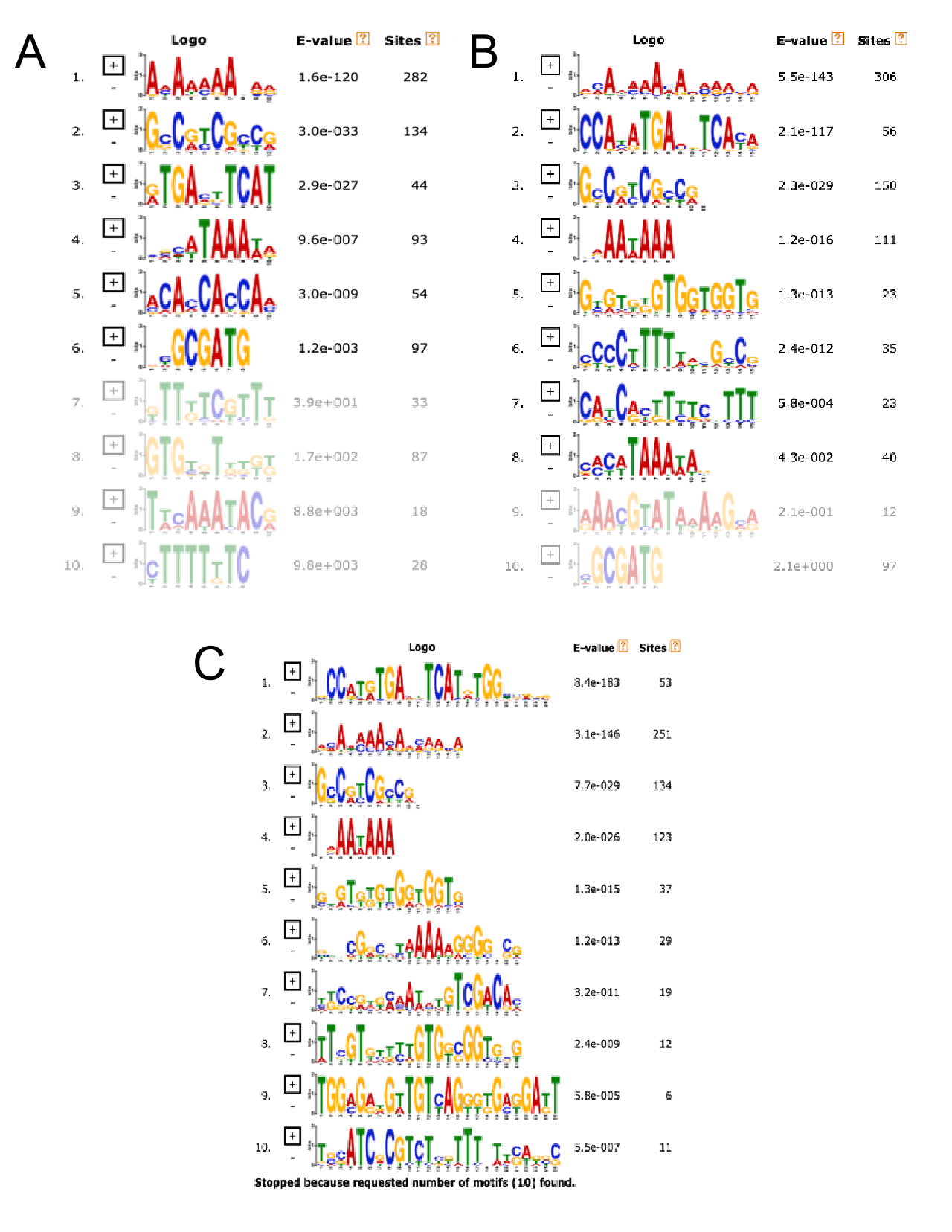

Fig.2 Promoter motif detected by MEME with motif width ranging from (A) 8-10, (B) 6-15, and (C) 8-25 bp. Motif sequences are ordered according to their e-value.

Table. 1 Motifs found under width parameter: 8-10 nucleotides

| Consensus sequence | width | sites | e-value |
| --- | --- | --- | --- |
| AMAAAAAVRR | 10 | 282 | 1.6e-120 |
| GSCRTCGCCG | 10 | 134 | 3.0e-033 |
| RTGAMTTCAT | 10 | 44 | 2.9e-027 |
| SVCATAAAWA | 10 | 93 | 9.6e-007 |
| ACACCACCAM | 10 | 54 | 3.0e-009 |
| DYGCGATG | 8 | 97 | 1.2e-003 |

Table. 2Motifs found under width parameter: 6-15 nucleotides

| Consensus sequence | width | sites | e-value |
| --- | --- | --- | --- |
| VMAAMAAMAVMAAMA | 15 | 306 | 5.5e-143 |
| CCAYATGAMBTCAYA | 15 | 56 | 2.1e-117 |
| GCCRYCGCCGH | 11 | 150 | 2.3e-029 |
| DRAAWAAA | 8 | 111 | 1.2e-016 |
| GYGTKKGTGGTGGTG | 15 | 23 | 1.3e-013 |
| CCCCTTTTWDHGYCG | 15 | 35 | 2.4e-012 |
| CAYCRYTTYTCDTTT | 15 | 23 | 5.8e-004 |

Table. 3 Motifs found under width parameter: 8-25 nucleotides

| Consensus sequence | width | sites | e-value |
| --- | --- | --- | --- |
| GCCATRTGAVKTCATRTGGYSRSG | 24 | 53 | 8.4e-183 |
| VMAAMAAMARMAAMA | 15 | 251 | 3.1e-146 |
| GCCRYCGYCGH | 11 | 134 | 7.7e-029 |
| NRAAWAAA | 8 | 123 | 2.0e-026 |
| GTGTKKGTGGTGGTG | 15 | 37 | 1.3e-015 |
| BBDCGRCRYWAAAAGGGGNSG | 21 | 29 | 1.2e-013 |
| YKCCRWKMAAWATGTCGACAC | 21 | 19 | 3.2e-011 |
