## Supplementary Material 6 for "RNA-seq of the medusavirus suggests remodeling of the host nuclear environment at an early infection stage"

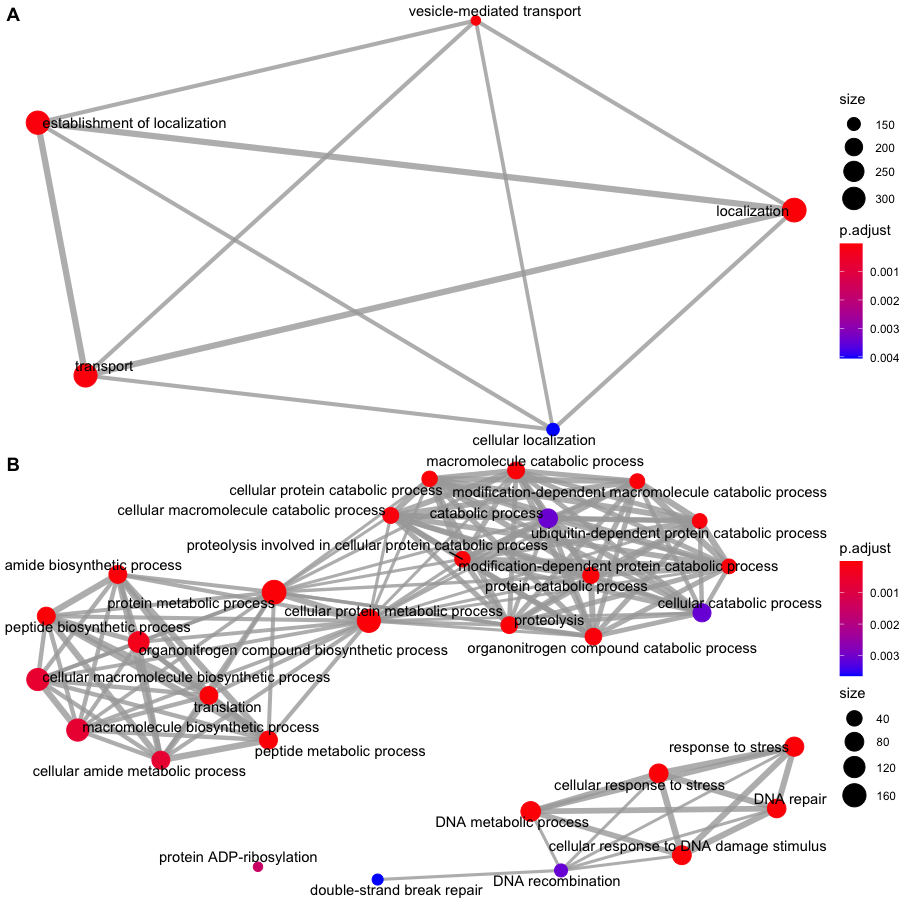

**Fig.1** Enriched map of enriched GO terms. A. GO terms in cluster 1; B. GO terms in cluster 2

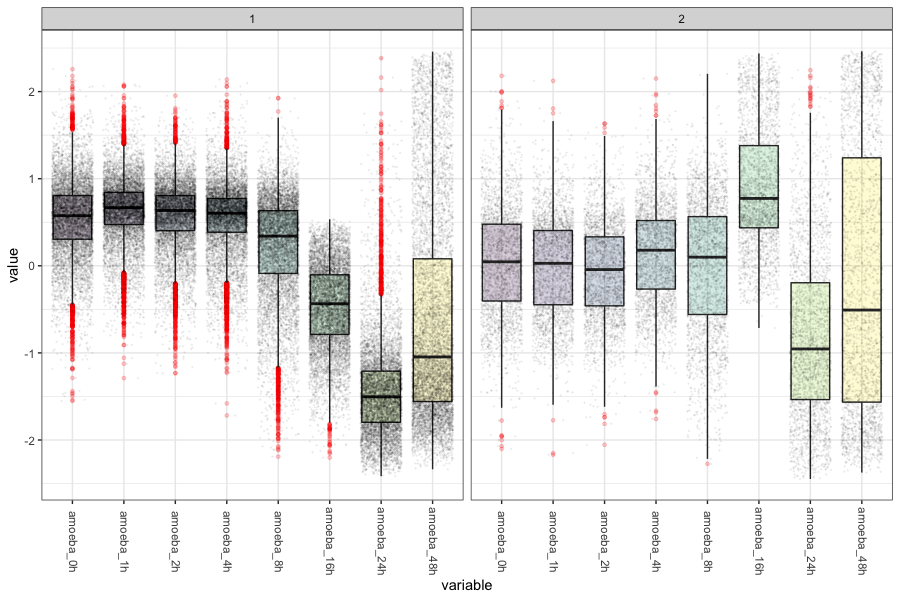

**Fig.2** Expression profile of 2 identified clusters at the end of the experiment.

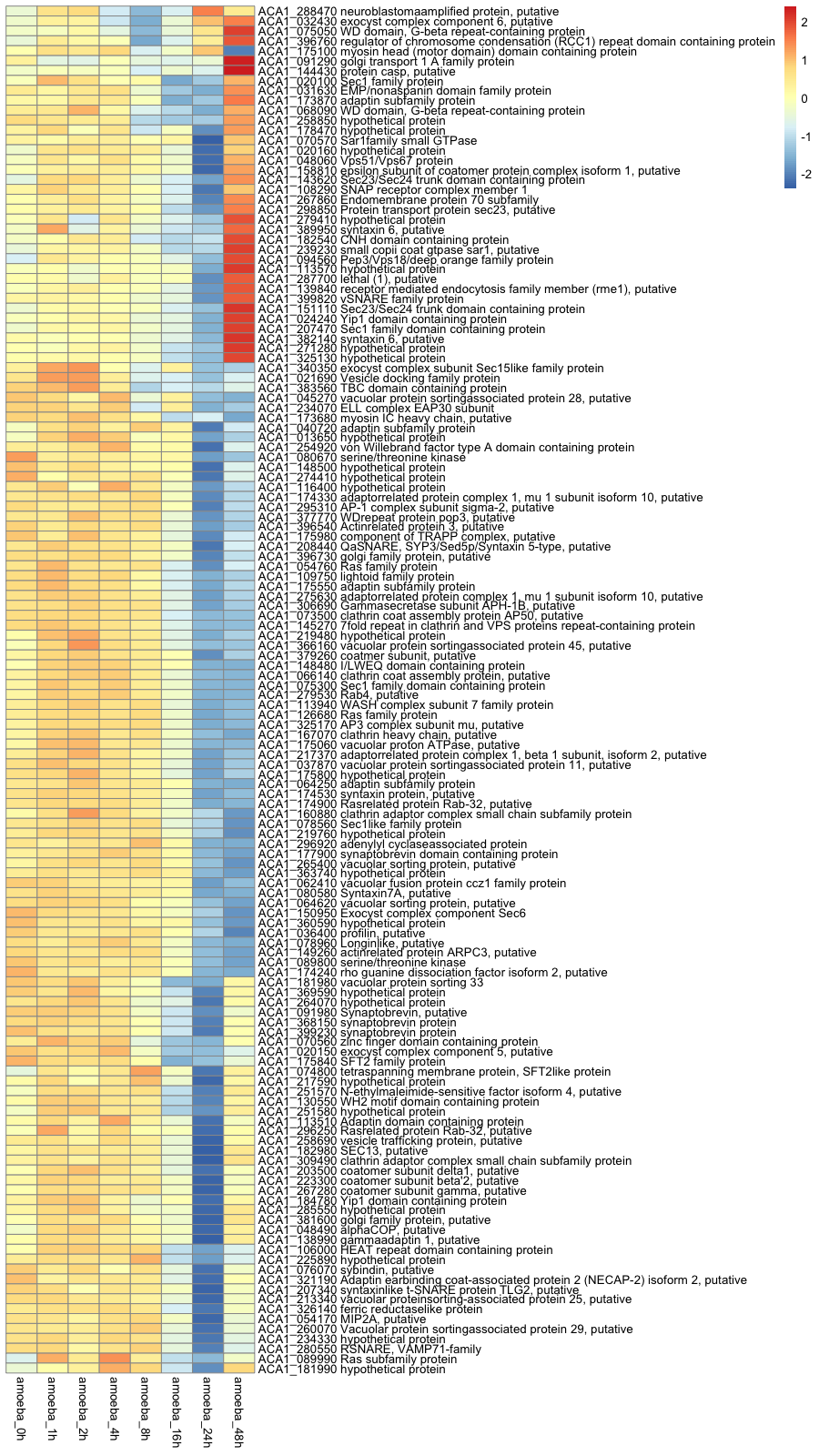

**Fig.3** Vesicle-mediated transport, GO:0016192;

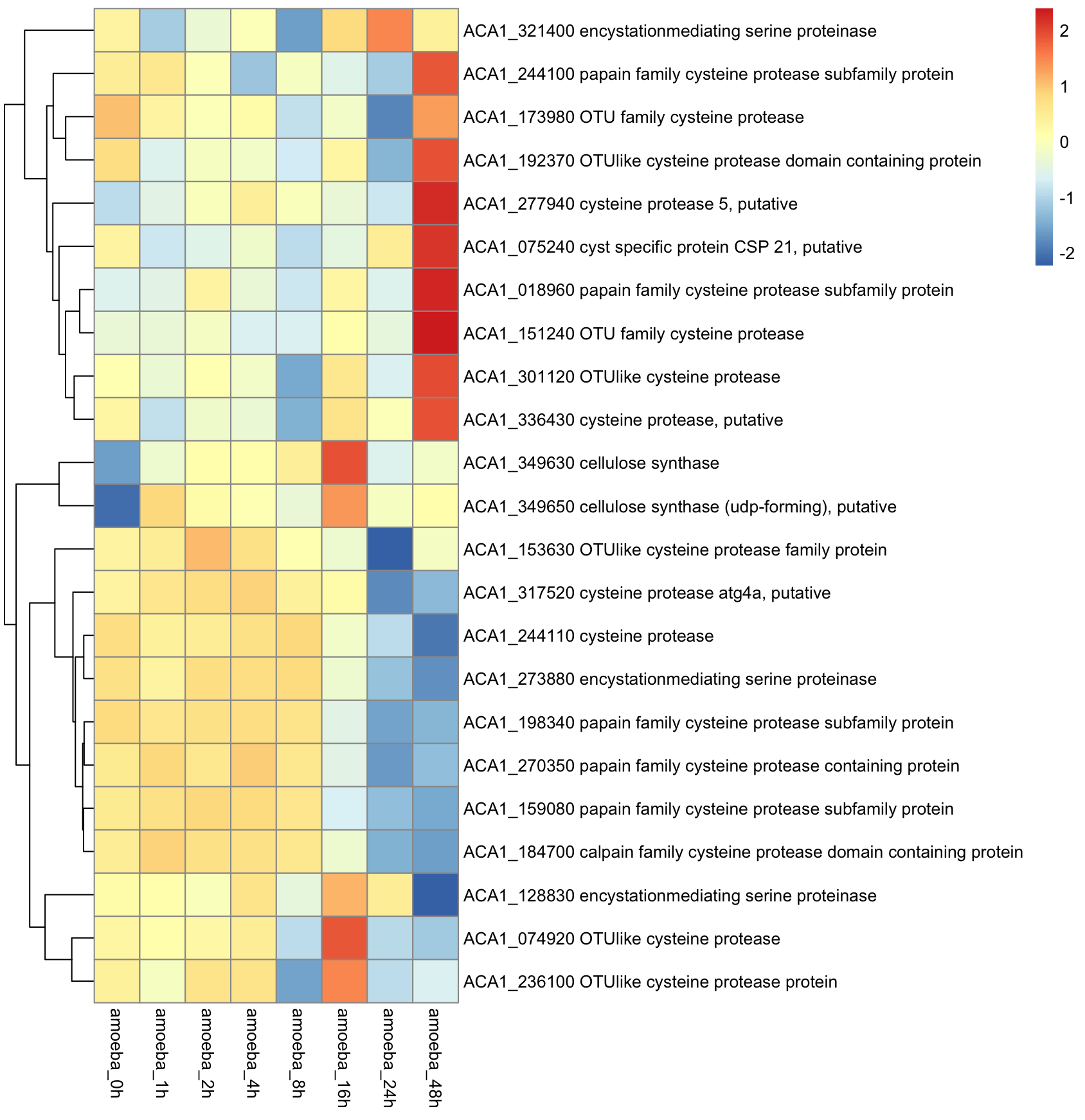

**Fig.4** Encystment related genes (gene name containing cysteine protease, CSP, encystation, or cellulose synthase)
